# Development of AlissAID system targeting GFP or mCherry fusion protein

**DOI:** 10.1101/2022.12.12.520189

**Authors:** Yoshitaka Ogawa, Kohei Nishimura, Keisuke Obara, Takumi Kamura

**Author notes:** Kohei Nishimura, Correspondence may also be addressed to Takumi Kamura. These two authors contributed equally to this work (Yoshitaka Ogawa, Kohei Nishimura).

## Abstract

Conditional control of target proteins using the auxin-inducible degron (AID) system provides a powerful tool for investigating protein function in eukaryotes. Here, we established an Affinity-linker based super-sensitive auxin-inducible degron (AlissAID) system in budding yeast by using a single domain antibody (a nanobody). In this system, target proteins fused with GFP or mCherry were degraded depending on a synthetic auxin, 5-Adamantyl-IAA (5-Ad-IAA). In AlissAID system, nanomolar concentration of 5-Ad-IAA induces target degradation, thus minimizing the side effects from chemical compounds. In addition, in AlissAID system, we observed few basal degradations which was observed in other AID systems including ssAID system. Furthermore, AlissAID based conditional knockdown cell lines are easily generated by using budding yeast GFP Clone Collection. From these advantages, the AlissAID system would be an ideal protein-knockdown system in budding yeast cells.

## INTRODUCTION

Conditional target-protein knockdown systems are used to analyze gene function. In particular, the auxin-inducible degron (AID) system(1-5) has been widely used to study protein-knockdown in various types of eukaryotes. In this method, AID-tagged target proteins are polyubiquitinated by SCF-OsTIR1 (Os referring to *Oryza sativa*), in an auxin-dependent manner. The polyubiquitinated proteins are then recognized and degraded by the 26S proteasome (6-9). The conventional AID system requires over 100 μM of indole-3-acetic acid (IAA) for target protein degradation, leading to cytotoxicity in some organisms (10).

To reduce the cytotoxicity caused by high IAA concentrations, super-sensitive AID (ssAID) (11) and auxin-inducible degron 2 (AID2) (12) systems have been developed in mammalian cell lines and applied fission yeast (13,14). Both systems utilize mutated OsTIR1 (the F74A mutant in ssAID and the F74G mutant in AID2) and enable target protein degradation by lowering the concentration of synthetic auxins. In the ssAID system with the OsTIR1^F74A^, a nanomolar concentration of 5-adamantyl-IAA (5-Ad-IAA) is sufficient for target-protein degradation via polyubiquitination. These improved AID systems could avoid cytotoxicity by reducing the concentration of inducer required for target protein degradation.

In these AID systems, an AID-tag must be fused with the target protein intended for degradation. Fusion with the AID-tag might affect target-protein stability and function. In some cases, co-expression with TIR1 causes weak degradation of the AID-tagged target without induction, which is termed basal degradation (15-17). One possible cause of basal degradation is that small amounts of IAA (or similar chemicals) induce degradation of target proteins. IAA is a metabolite of tryptophan, and similar metabolites are present in non-plant organisms as well (18,19). Owing to basal degradation, it is sometimes difficult to generate AID-based conditional knockdown cell lines for essential proteins. Although some improved AID systems (including ssAID and AID2) showing limited basal degradation have been developed in mammalian cell lines(11,12,16,17) and fission yeast (13), It has been reported that OsTIR1^F74A^ showed a mild sensitivity to IAA at 30 °C in budding yeast (11).

Single-peptide antibody nanobodies (20,21) are useful for protein recognition in target protein degradation systems. Several studies have reported that fusion proteins of E3 ligase and the nanobody can induce degradation of target proteins recognized by nanobodies. Such targeted degradation systems have been developed in studies using *Drosophila* (22) and have been applied to human cells (23-25). The VHHGFP4 nanobody, which binds to green fluorescent protein (GFP) has been applied to the conventional AID system (26). GFP fusion protein was degraded in cultured human and zebrafish cells expressing both wild-type OsTIR1 (OsTIR1^WT^) and a chimeric protein-fused AID-tag containing VHHGFP4. In this chimeric protein, the lysine residues have been replaced by arginine residues; as a result, OsTIR1 can ubiquitinate the target protein bound to the chimeric protein without ubiquitinating the chimeric protein. The GFP-tag is a stable tag that is widely used for target-protein visualization, and GFP-tagged lines have been generated for various species. Thus, A GFP fusion-protein-targeting AID system represents a powerful molecular research tool for eukaryotes.

The budding yeast *Saccharomyces cerevisiae* is a model organism for eukaryotic cells. In budding yeast, the GFP-tag can be easily fused with an endogenous protein via genetic manipulation. A Yeast GFP Clone Collection (27) (Thermo Fisher scientific), with a GFP-tagged Open Read Frame (ORF) at its chromosomal locus, that contains 4,159 GFP-tagged ORFs, comprising 75% of the yeast proteome, is currently available. We combined a nanobody system with our ssAID system to develop an Affinity-linker based super-sensitive auxin-inducible degron (AlissAID) system in budding yeast. AlissAID-based conditional-knockdown cells can be generated by the expression both OsTIR1^F74A^ and a chimeric protein

AID-tag fused with the nanobody in the GFP-tagged strain. This system utilizes 5-Ad-IAA as a degradation inducer and induces target-protein degradation at nanomolar concentrations of inducer. We also found that it can be applied to the degradation of mCherry fusion proteins, which is recognized with LaM2 or LaM4 nanobodies. The AlissAID system is therefore useful to analyze target-protein physiological function in budding yeast cells.

## MATERIAL AND METHODS

### Yeast strains and media

The *Saccharomyces cerevisiae* strains used in this work are listed in Supplementary Table 1. Cells were grown in YPD (2% D-glucose, 1% yeast extract, and 2% peptone,) supplemented with 100 mg/L adenine or SD medium (2% D-glucose and 0.67% yeast nitrogen base without amino acids) supplemented with appropriate amino acids, uracil, tryptophan, histidine, and leucine.

### Auxin derivative stock solutions

3-Indoleacetic Acid (IAA) (Nacalai tesque: 19119-61) and 5-Adamantyl-IAA (Tokyo Chemical Industry: A3390) were diluted into dimethyl sulfoxide (DMSO) at 500 mM or 5 mM concentration and stored at -30 °C. When mixed with plate medium, a solution of 1000 times the concentration of each final concentration was prepared, and the medium was prepared by adding 1/1000 to the medium.

### Genetic manipulation

Chromosome fusions of *GFP, mAID-GFP* and *mCherry* to the 3′-terminus of the gene were conducted using PCR-based gene modification (28). The sequence containing the tag [*GFP, mAID-GFP, mCherry*], the *ADH1* terminator, and a marker gene was amplified using PCR from pFA6a GFP-His3MX6 [for GFP], pFA6a mAID-GFP-His3MX [for mAID-GFP](pYS15, this study), pFA6a mCherry-His3MX6[for mCherry](our laboratory) with a primer set containing the homologous region(50-55 mer)of each gene. The PCR-amplified fragments were directly inserted into the chromosome via homologous recombination. Successful deletions of the genes and tagging were confirmed via genomic PCR, immunoblot, and/or fluorescence microscopy. The sequence encoding *OsTIR1(F74A), OsTIR1(F74G), mAID(KR)-VHHGFP4(KR), mAID(KR)-LaM2(KR), mAID(KR)-LaM4(KR), mAID(KR)-LaM8(KR)* is integrated into *URA3* or *LEU2* locus as follows. The *ADH1* promoter sequence followed by *OsTIR1(F74A), OsTIR1(F74G), mAID(KR)-VHHGFP4(KR), mAID(KR)-LaM2(KR), mAID(KR)-LaM4(KR), mAID(KR)-LaM8(KR)* sequence was constructed on pRS306 or pRS305 (29). The resultant plasmids were linearized by digestion with *Stu*1 or *Bst*X1 or *Afl*2 and integrated into *URA3* or *LEU2* locus, respectively, via homologous recombination. The sequence encoding *OsTIR1(WT)-T2A-mAID(KR)-VHHGFP4(KR), OsTIR1(F74A)-T2A-mAID(KR)-VHHGFP4(KR), OsTIR1(F74S)-T2A-mAID(KR)-VHHGFP4(KR), OsTIR1(F74C)-T2A-mAID(KR)-VHHGFP4(KR)* is integrated into *URA3* locus as follows. The *ADH1* promoter sequence followed by *OsTIR1(WT)-T2A-mAID(KR)-VHHGFP4(KR), OsTIR1(F74A)-T2A-mAID(KR)-VHHGFP4(KR), OsTIR1(F74S)-T2A-mAID(KR)-VHHGFP4(KR), OsTIR1(F74C)-T2A-mAID(KR)-VHHGFP4(KR)* was constructed on pRS306. And BY4741 *URA3* locus sequence is constructed between *Eco*R1 and *Xho*1 site on same plasmids. The resultant plasmids were linearized by digestion with *Sma*1 and integrated into the *URA3* locus, respectively, via homologous recombination.

### Immunoblot analysis

Cell lysates were prepared using the alkaline-trichloroacetic acid method. Harvested cells were resuspended in an ice-cold alkaline solution containing 0.25 M of NaOH and 1% (v/v) of 2-mercaptoethanol. After incubation on ice for 10 min, Trichloroacetic acid was added at a final concentration of 7% (w/v). After centrifugation at 20,000 × g for 2 min at 4 °C, the supernatant was removed. Then, 1 M tris was gently added, ensuring to not break the pellet, to neutralize the remaining trichloroacetic acid, and centrifuged at 20,000 × g for 30 s. After the removal of supernatant, proteins were eluted by incubation in SDS-sample buffer for 5 min at 95 °C. After centrifugation at 10,000 × g for 1 min at room temperature, the supernatant was transferred to a new tube and subjected to immunoblotting. Proteins were separated via SDS-PAGE and transferred to an ImmobilonTM polyvinylidene difluoride membrane (Millipore, Billerica, USA). The membrane was incubated with anti-GFP (1:2000 dilution; our laboratory), anti-RFP(1:2000 dilution; Chromotek, 5f8-20/5f8-100), anti-Pgk1 (1:5000 dilution; our laboratory), anti-TIR1 (1:2000 dilution; Medical and Biological Laboratories, PD048), anti-AID (1:2000 dilution; gifted from Prof. Karim Labib). HRP-conjugated anti-Rabbit IgG (1:5000 dilution; Sigma, A6154) or Peroxidase-conjugated anti-Sheep IgG (1:5000 dilution; 713-035-003, Jackson ImmnoResearch) or Peroxidase-conjugated anti-Rat IgG(1:5000 dilution; 112-035-003, Jackson ImmnoResearch) was used as the secondary antibody. Immunodetection was performed using a Luminata Forte Western HRP Substrate system (Merck Millipore, Burlington, USA, 61-0206-81) or a Chemi-Lumi One L system (Nacalai Tesque, 07880) with a bioanalyzer (LAS4000 mini; GE Healthcare Biosciences).

### Fluorescence microscopy

Cells were grown to log phase in SD medium containing 5 μM 5-Ad-IAA for 3 hours at 30 °C. After centrifugation, cells were fixed by adding 4% formaldehyde in PBS for 10 min at RT. Cells were collected by centrifugation. After washing with PBS, cells were resuspended in PBS and observed under a fluorescence microscope (AxioObserver Z1; Carl Zeiss, Oberkochen, Germany) equipped with a CCD camera (AxioCam MRm; Carl Zeiss). Exposure time GFP:2,000 ms, mCherry:4000 ms. Fluorescence intensity was analyzed using ImageJ for photographs taken under the same conditions.

### Spot assay (serial dilution assay)

Yeast cells were grown in YPD or SD medium overnight at 30°C until OD_600_ reached 0.7-0.8. Cells were collected and suspended in 200 μL of sterile water (0.3 OD_600_ equivalent). Ten-fold serial dilutions were generated as follows. 200 μL of the cell suspension was transferred to the first lane of a 96-well plate. 180 μL of sterile water were placed in the second, third, and fourth lanes. Then, 20 μL of the cell suspension in the first lane was transferred to the second lane (lane 2) and mixed by pipetting ten times. Similarly, 20 μL of the cell suspension was transferred from the second lane to the third lane, and from the third lane to the fourth lane, as described above. An aliquot of the cell suspension (3 μL on YPD plate, 5 μL on SD plate) was spotted on agar plates containing IAA, 5-Ad-IAA, or G418, which were then incubated at 30°C.

### Flow cytometry

Cells were grown in YPD medium overnight at 25°C not to exceed OD_600_ =1. After treated with 5μM 5-Ad-IAA at 30°C, cells were centrifugated (3000 g, 5 min) to pellet and discard the medium. Cells were resuspended and fixed in 70% ethanol at -30°C for 1Day. Fixed cell were centrifugated, treated with 100 μl stain mix (PBS, 1mg/ml RNase, 1/1000 SYBR™ Green I Nucleic Acid Gel Stain, Thermo Fisher Scientific) at 37°C for 30 min. Add 900 μl PBS, vortex, cells were acquired by Attune Flow Cytometer (Thermo Fisher Scientific).

## RESULTS

### GFP fusion protein is rapidly degraded in OsTIR1^F74A^ and mAID-Nb expressing cell lines

The AID system enables rapid degradation of AID-tagged target proteins within cells via the natural auxin IAA, requiring approximately 500 μM of IAA for target protein degradation (Figure 1A) (1). In contrast, the ssAID system, which uses the OsTIR1^F74A^ mutant and an artificial auxin, 5-Ad-IAA, allows target-protein degradation at a nanomolar concentration of 5-Ad-IAA (Figure 1B) (11). Therefore, by using the ssAID system, it may be possible to avoid the side effects of high concentrations of degradation inducers. In both AID systems, AID-tags are necessary for target protein degradation. However, AID-tags can affect target-protein stability or function. Here, we attempted to eliminate these effects by using nanobodies for target-protein recognition (Figure 1C). In the AlissAID system, the AID-tagged nanobody binds to the target protein. The AID-tag was recruited to OsTIR1^F74A^ in a 5-Ad-IAA-dependent manner. All the lysine residues of the AID-tagged nanobody were replaced with arginine, preventing it from being ubiquitinated by the SCF complex and OsTIR1^F74A^. Instead, the target protein is ubiquitinated and degraded by the 26S proteasome.

**Figure 1.**
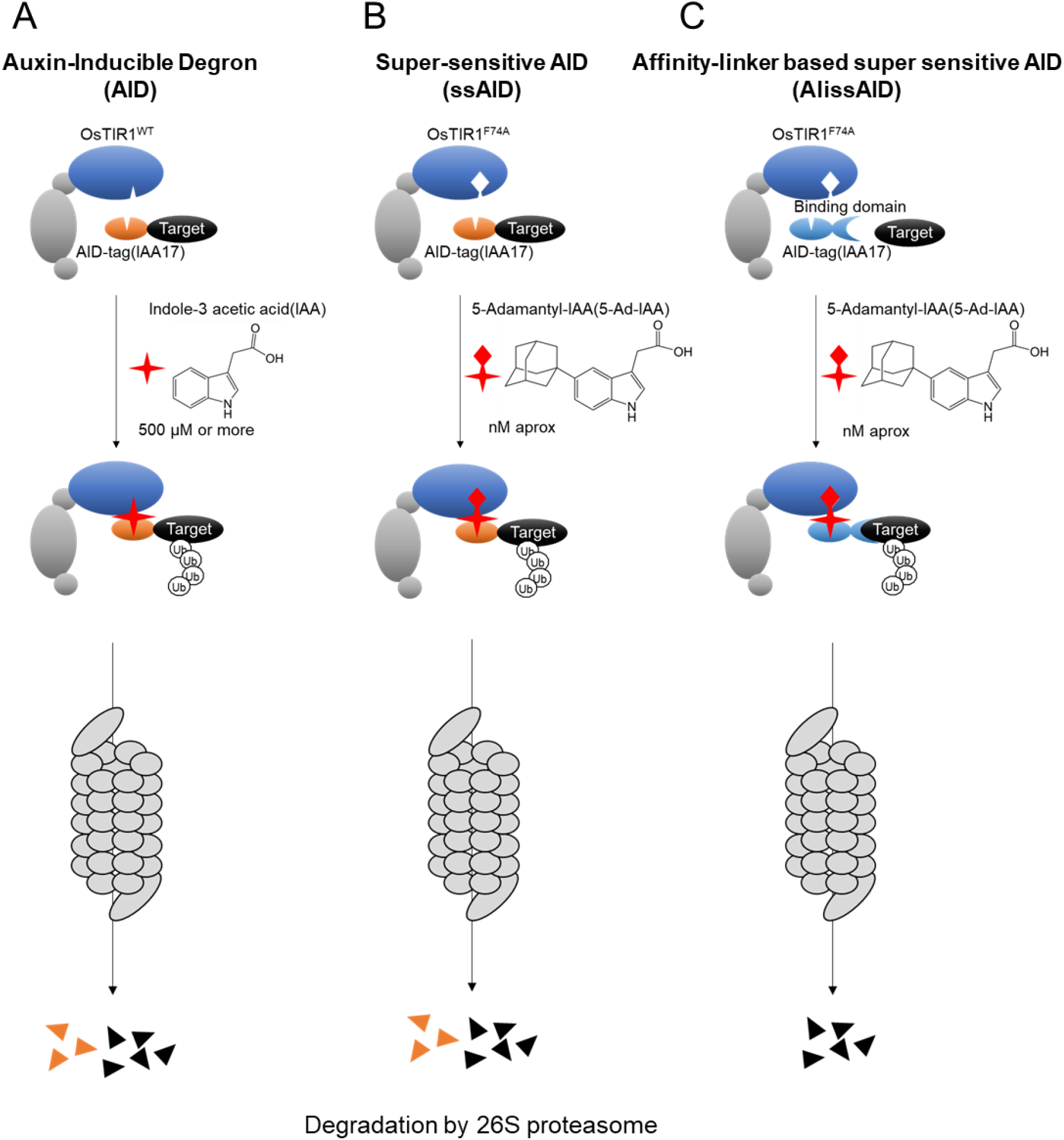
Schematic illustration of (A) the Auxin-Inducible Degron (AID), (B) super-sensitive AID (ssAID), and (C) Affinity-linker based super sensitive AID (AlissAID) systems.

We attempted to establish an AlissAID system in *S. cerevisiae*. We expressed both OsTIR1^F74A^ and a minimal AID-tag fused with an anti-GFP nanobody (mAID-Nb), in which all lysine residues were replaced by arginine residues, under the control of the constitutive ADH1 promoter in the W303-1a background (30). We added a GFP-tag to the essential proteins Mcm4, Ask1, and Neo1. Immunoblotting revealed that target proteins were degraded within 1 h of 5-Ad-IAA addition (Figure 2A). Fluorescent microscopy revealed that, following 5-Ad-IAA addition, Mcm4-GFP (localized in the nucleus and cytoplasm), Ask1-GFP (localized to the nucleus), and Neo1-GFP (localized at the membrane) GFP signals were reduced (Figure 2B). For these cell lines expressing OsTIR1^F74A^ and mAID-Nb, serial dilution spotting to confirm growth-defect phenotypes revealed severe growth defects, even in the presence of 50 nM 5-Ad-IAA (Figure 2C). Mcm4 and Ask1 have been reported to be essential for the progression of G1/S and G2/M phases (31,32), respectively. FACS analysis showed that Mcm4-GFP cells were arrested at G1/S phase and Ask1-GFP cells were arrested at G2/M phase 1 h after the addition of 5-Ad-IAA in the AlissAID system (Supplementary Figure 1A, B). These results indicate that the AlissAID system works well in budding yeast by enabling the degradation of nuclear, cytosolic, and membrane proteins at low concentrations of 5-Ad-IAA.

**Figure 2.**
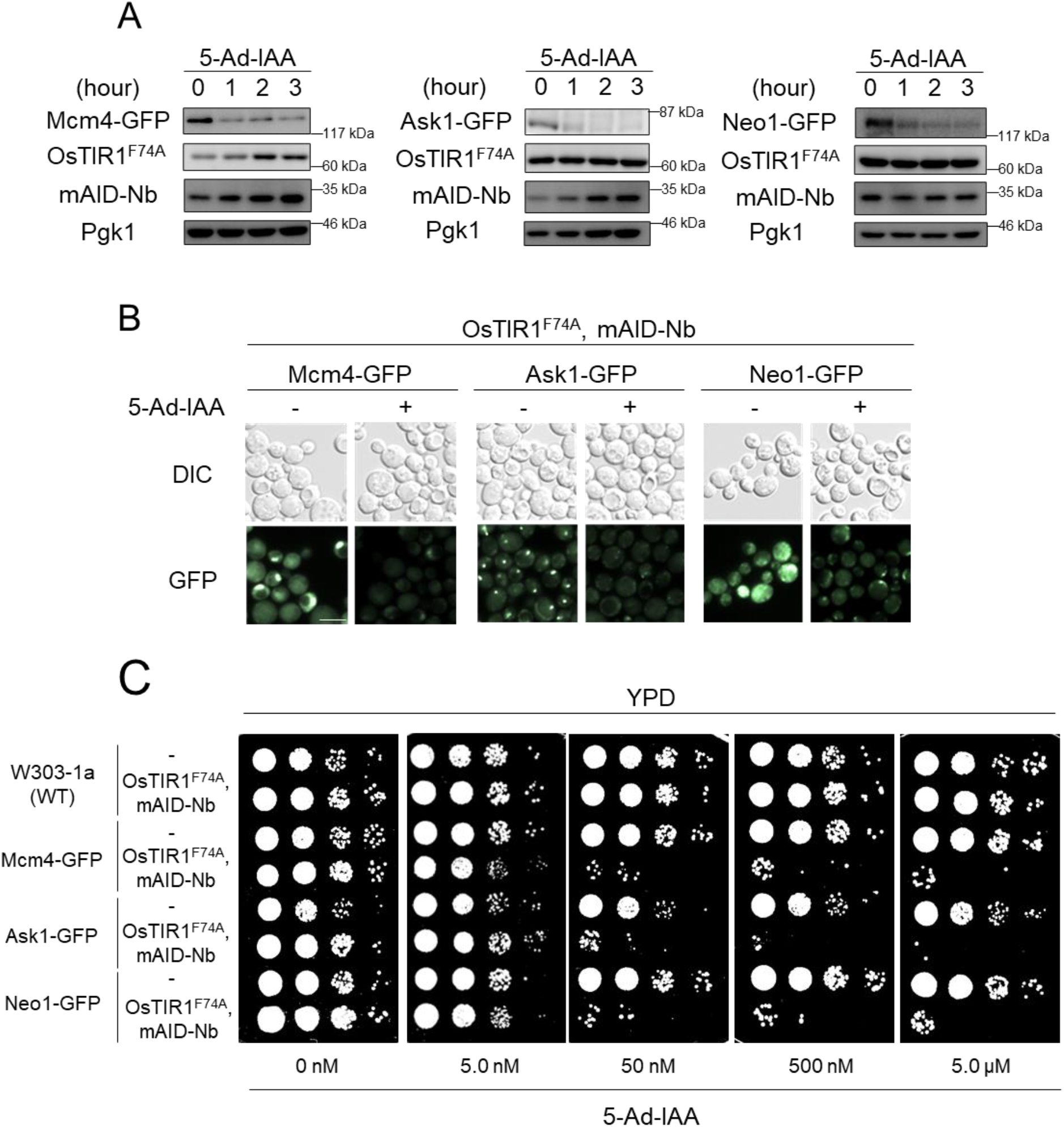
Efficient degradation of GFP fusion proteins in OsTIR1^F74A^ and mAID-Nb-expressing cells in the presence of 5-Ad-IAA. (A) Immunoblotting analysis of GFP fusion proteins Mcm4, Ask1, and Neo1. Cells were treated with 5 μM 5-Ad-IAA and sampled at the indicated time points. Pgk1 was used as a loading control. (B) Fluorescence microscopy observations of Mcm4-GFP, Ask1-GFP, or Neo1-GFP. Cells are treated with or without 5.0 μM 5-Ad-IAA for 3 h. White bar, 5 μm. (C) Serial dilution spotting on YPD medium containing 5-Ad-IAA. Cells were grown for 30 h at 30 °C.

### AlissAID system reduce target-protein basal degradation

Basal degradation is among the main drawbacks of the AID system. Some proteins are degraded even in the absence of auxin in the conventional AID system(15-17). Minor amounts of auxin (natural auxin), a metabolite of tryptophan, are present even in non-plant organisms, which may induce the binding of TIR1 and AID-tag. Several reports showed that basal degradation rarely occurs in ssAID because the natural auxin does not affect OsTIR1^F74A^ (11,12). We coincidentally found that mAID-taged Neo1 proteins is reduced by basal degradation in budding yeast ssAID system. There are questions about the extent to which basal degradation occurs in the AlissAID system. To approach this, we evaluated basal degradation in the ssAID and AlissAID systems. For the ssAID system, we generated a Neo1-conditional-knockdown cell line by tagging mAID-GFP at the C-terminus of Neo1 in OsTIR1^F74A^-expressing cells. For the AlissAID system, we generated a Neo1-conditional-knockdown cell line by tagging GFP at the C-terminus of Neo1 in both OsTIR1^F74A^- and mAID-Nb-expressing cells. After comparing Neo1-mAID-GFP in the ssAID system with Neo1-GFP in the AlissAID system, we observed a substantial reduction in Neo1 protein levels even in the absence of 5-Ad-IAA in the ssAID system (Figure 3A,B). In addition, Neo1-mAID-GFP levels were found to be reduced by co-expression with OsTIR1^F74A^, while Neo1-GFP levels were not affected by co-expression with OsTIR1^F74A^ and mAID-Nb (Figure 3C). Mcm4-GFP and Ask1-GFP levels are also not reduced by co-expression of OsTIR^F74A^ and mAID-Nb (Supplemental Figure 2A). As Neo1 protein is associated with G418 resistance in budding yeast, we compared the G418 sensitivity of the two strains via serial dilution spotting in the absence of 5-Ad-IAA. The ssAID-cells exhibited defective growth on G418-containing medium, whereas the AlissAID-cells did not show any growth defects under the same condition (Figure 3D). This indicates that the AlissAID system shows milder basal degradation levels than the ssAID system. Furthermore, minimal target degradation was observed in AlissAID strains with OsTIR1^F74G^, which has been reported to reduce basal degradation. (Supplementary Figure 2B, C). These results suggest that the AlissAID system with OsTIR1^F74A^ would be a good candidate for a low-basal-degradation system in budding yeast.

**Figure 3.**
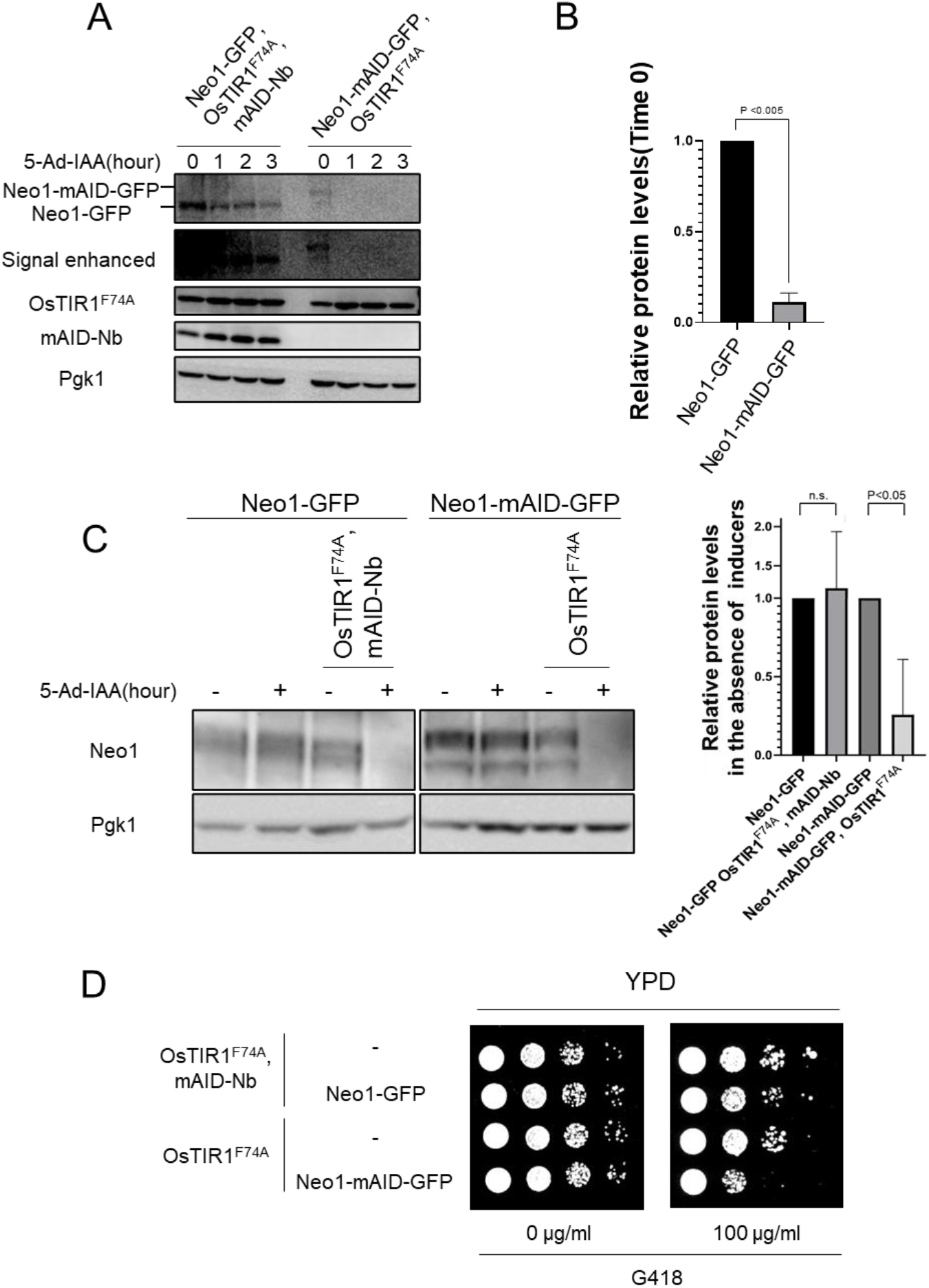
The AlissAID system reduces basal degradation and side effects. (A) Immunoblotting of endogenous tagged Neo1-GFP and Neo1-mAID-GFP. Cells were treated with 5 μM 5-Ad-IAA and sampled at the indicated time points. Pgk1 was used as a loading control. (B) Protein levels relative to those at time zero, normalized to those of Pgk1 (based on three independent experiments.) (C) Immunoblotting of Neo1-GFP and Neo1-mAID-GFP. Cells were treated with or without 5 μM 5-Ad-IAA for 3 h. Protein levels relative to the level in OsTIR1^F74A^ and mAID-Nb (AlissAID), OsTIR1^F74A^ (ssAID) un-expression cell, normalized to those of Pgk1 (based on three independent experiments). (D) Serial dilution spotting on YPD medium containing G418 (Geneticin) and grown for 24 or 54 h at 30 °C.

### Use of a yeast GFP Clone Collection in conjunction with the AlissAID system enables easy generation of a degron cell line

There is the budding yeast GFP Clone Collection (27) in the BY4741(33) background. The GFP gene is integrated into the C-terminus of the endogenous gene via homologous recombination, and the GFP fusion protein is expressed under the endogenous promoter. This fusion collection is useful for generating GFP-degron cells via the AlissAID system to target budding yeast proteins (Figure 4A). Therefore, we constructed a plasmid by combining OsTIR1^F74A^ and mAID-Nb under the control of the *ADH1* promoter on pRS316 (29), with OsTIR1^F74A^ and mAID-Nb connected by self-cleavage T2A sequence (Figure 4B). First, we generated the Neo1-degron strain by transforming the plasmid into Neo1-GFP. Neo1-GFP protein was rapidly degraded after the addition of 5-Ad-IAA (Figure 4C). Subsequently, we generated an AlissAID strain with various essential proteins by transforming this plasmid into GFP clones. While all the cells grew well in the absence of 5-Ad-IAA, severe growth defects were observed in the presence of 500 nM 5-Ad-IAA (Figure 4D), and GFP signals were reduced in the presence of 5-Ad-IAA (Supplementary Figure 3). These results indicate that the combined application of a GFP clone collection and the AlissAID system enables easy generation of degron cell lines. This combined system offers great potential for studying protein function in budding yeast.

**Figure 4.**
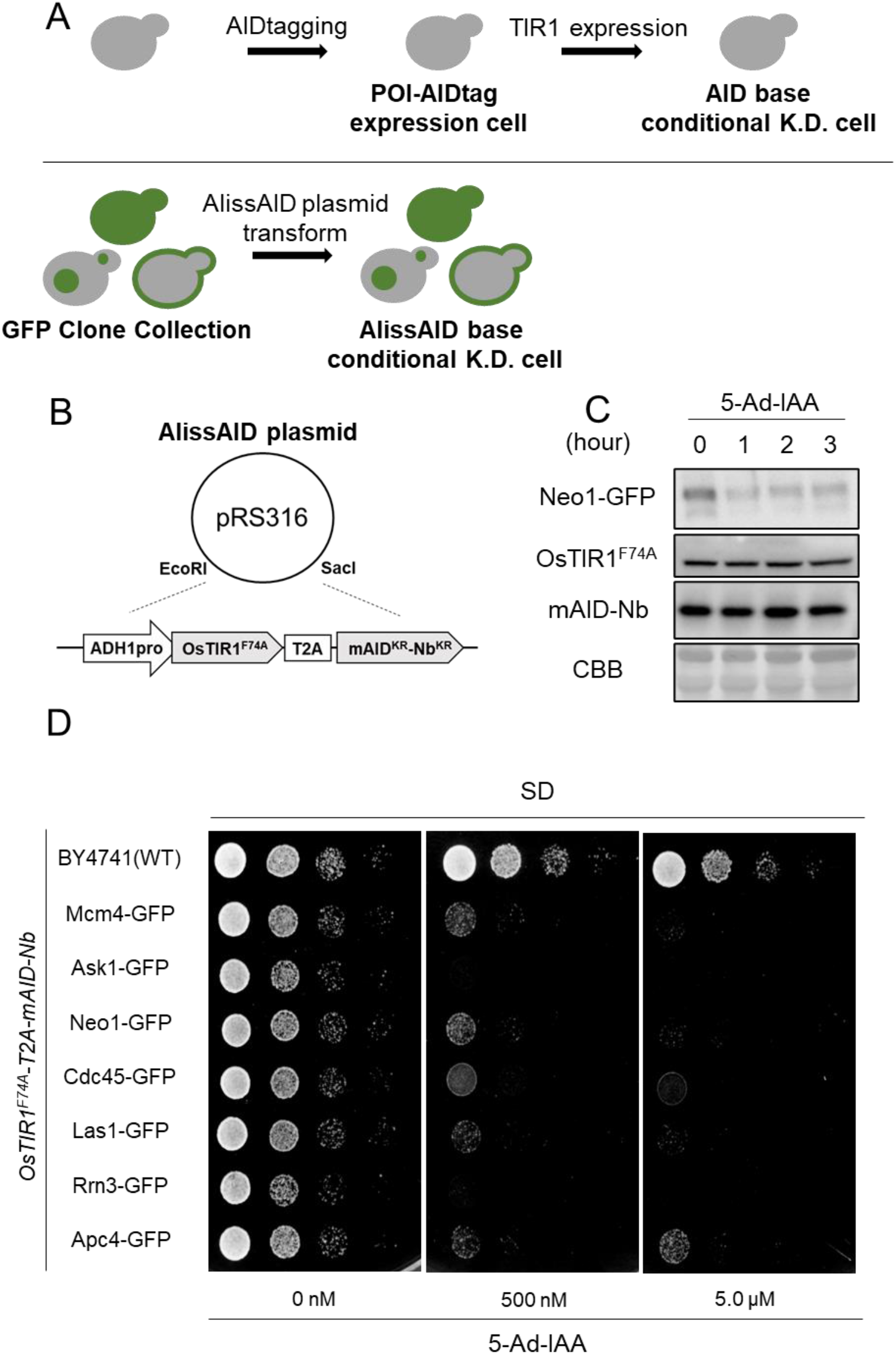
Target-protein degradation and phenotyping in GFP Clone cells with single-plasmid coding OsTIR1^F74A^ and mAID-Nb. (A, B) Degron strain preparation and plasmid construct. (C) Immunoblotting of endogenous GFP-tagged Neo1 cells transformed with the plasmid. Cells were treated with 5 μM 5-Ad-IAA. (B) Serial dilution spotting on SD medium containing 5-Ad-IAA. Cells were grown for 24 h at 30 °C.

### OsTIR1^F74A^ is suitable for the AlissAID system

The use of OsTIR1^WT^ with nanobodies can degrade GFP fusion protein in mammalian cells and zebrafish embryos (26). Additionally, in mammalian cell lines, certain OsTIR1 mutants effectively degrade target proteins (11). To compare the degradation effectiveness of OsTIR1 mutants, we generated an Mcm4-GFP degron strain by expressing OsTIR1 (wild-type [WT], and mutants F74A, F74S, or F74C) with mAID-Nb. Mcm4-GFP was severely degraded in the presence of 5 μM 5-Ad-IAA in OsTIR1^F74A^, while it was only slightly degraded in the OsTIR1^F74S^ and OsTIR1^F74C^ strains. However, Mcm4-GFP was barely degraded following the addition of 500 μM IAA in the OsTIR1^WT^ strain (Figure 5A, B). A similar result was obtained via serial dilution spotting analysis. OsTIR1^F74A^-expressing cells exhibited severe growth defects, even in the presence of 50 nM 5-Ad-IAA. OsTIR1^F74S^- and OsTIR1^F74C^-expressing cells required 500 nM 5-Ad-IAA to generate growth defects, while OsTIR1^WT^-expressing cells required 500 μM IAA (Figure 5C). These results indicate that OsTIR1^F74A^ is the suitable OsTIR1 mutant for use in the AlissAID system in budding yeast.

**Figure 5.**
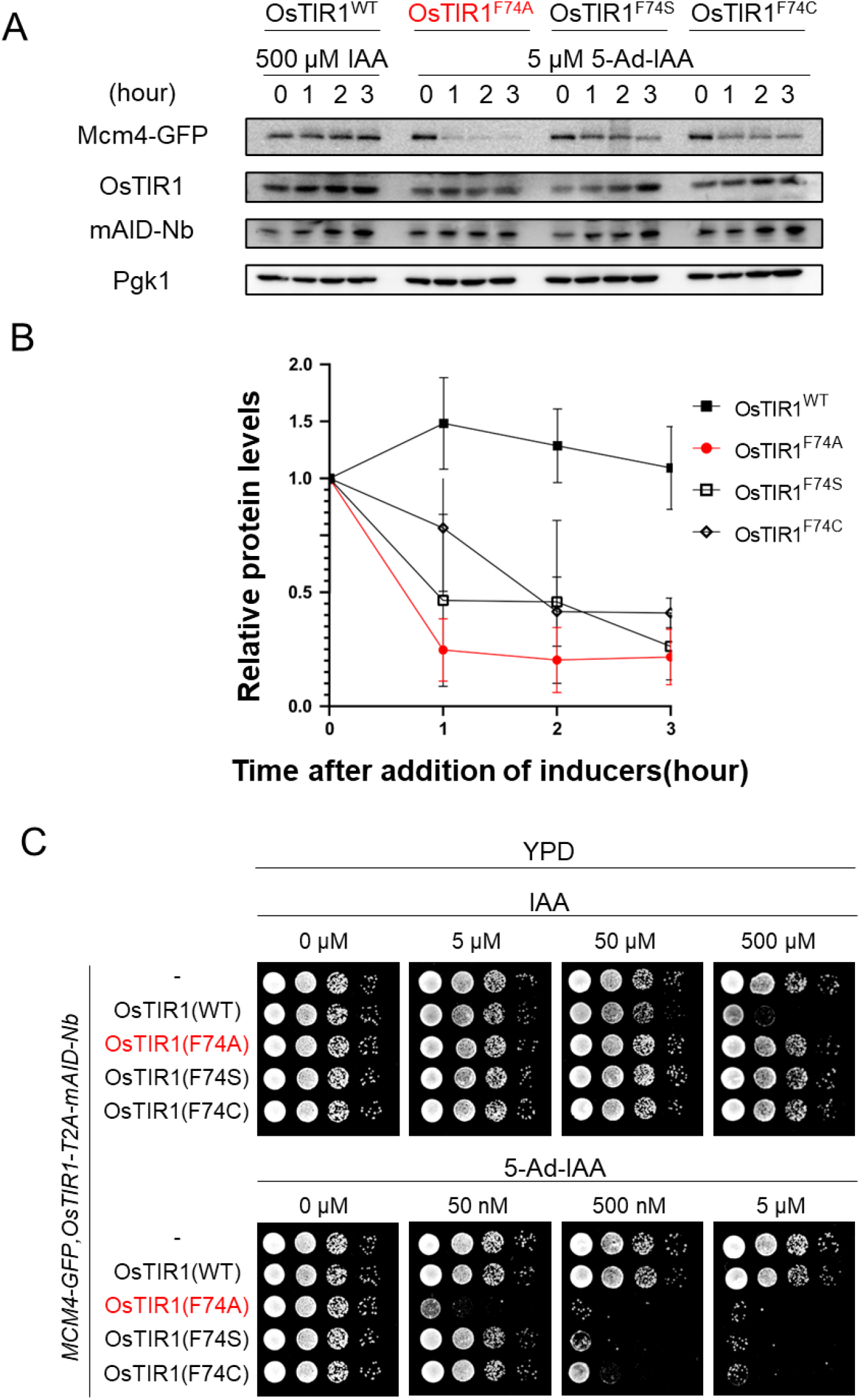
The OsTIR1^F74A^ mutant most efficiently removes target proteins. (A, B) Immunoblotting of endogenous tagged Mcm4-GFP cells treated with 500 μM IAA or 5 μM 5-Ad-IAA and sampled at the indicated time points. Pgk1 was used as a loading control. Protein levels are relative to those at time zero and normalized to those of Pgk1 (based on three independent experiments). (C) Serial dilution spotting on YPD medium containing IAA or 5-Ad-IAA. Cells were grown for 24 h at 30 °C.

### mCherry fusion proteins were degraded by the AlissAID system using anti-mCherry nanobody, LaM2 and LaM4

GFP is among the most popular fluorescent proteins used in budding yeast. Our AlissAID system for GFP fusion protein utilizes OsTIR1^F74A^ and a mAID-tagged nanobody that recognizes GFP. mCherry is commonly used as a red fluorescent protein in budding yeast. We therefore attempted to generate a conditional knockdown cell line using the AlissAID system with anti-mCherry nanobodies for mCherry fusion proteins. We added mCherry tag to the C-terminus of the essential protein Ask1 via homologous recombination and expressed both OsTIR1^F74A^ and mAID-LaM2 and LaM4 or LaM8 (34) in the W303-1a background (Supplementary Figure 4A). These cells expressed the same levels of each mAID-Nb (LaM2, LaM4, or LaM8) (Supplementary Figure 4B). Immunoblotting analysis revealed that Ask1-mCherry was highly degraded in cell lines with mAID-LaM2 and weakly degraded in cell lines with mAID-LaM4, but not in cell lines with mAID-LaM8 (Figure 6A). Further, after the addition of 5-Ad-IAA, Ask1-mCherry fluorescence signals were diminished in mAID-LaM2- and mAID-LaM4-expressing cells but not in mAID-LaM8-expressing cells (Figure 6B, Supplementary Figure 4C). The mAID-LaM2 expressing cell line showed severe growth defects in the presence of 5-Ad-IAA in serial dilution spotting assay. While the mAID-LaM4 expressing cell line showed mild growth defects, the mAID-LaM8 expressing cell line did not show any growth defects (Figure 6C). A similar trend was also observed in degron strains targeting Neo1-mCherry-Flag (Supplementary Figure 4D). These results indicate that LaM2 is a good candidate for the AlissAID system for mCherry fusion proteins in budding yeast.

**Figure 6.**
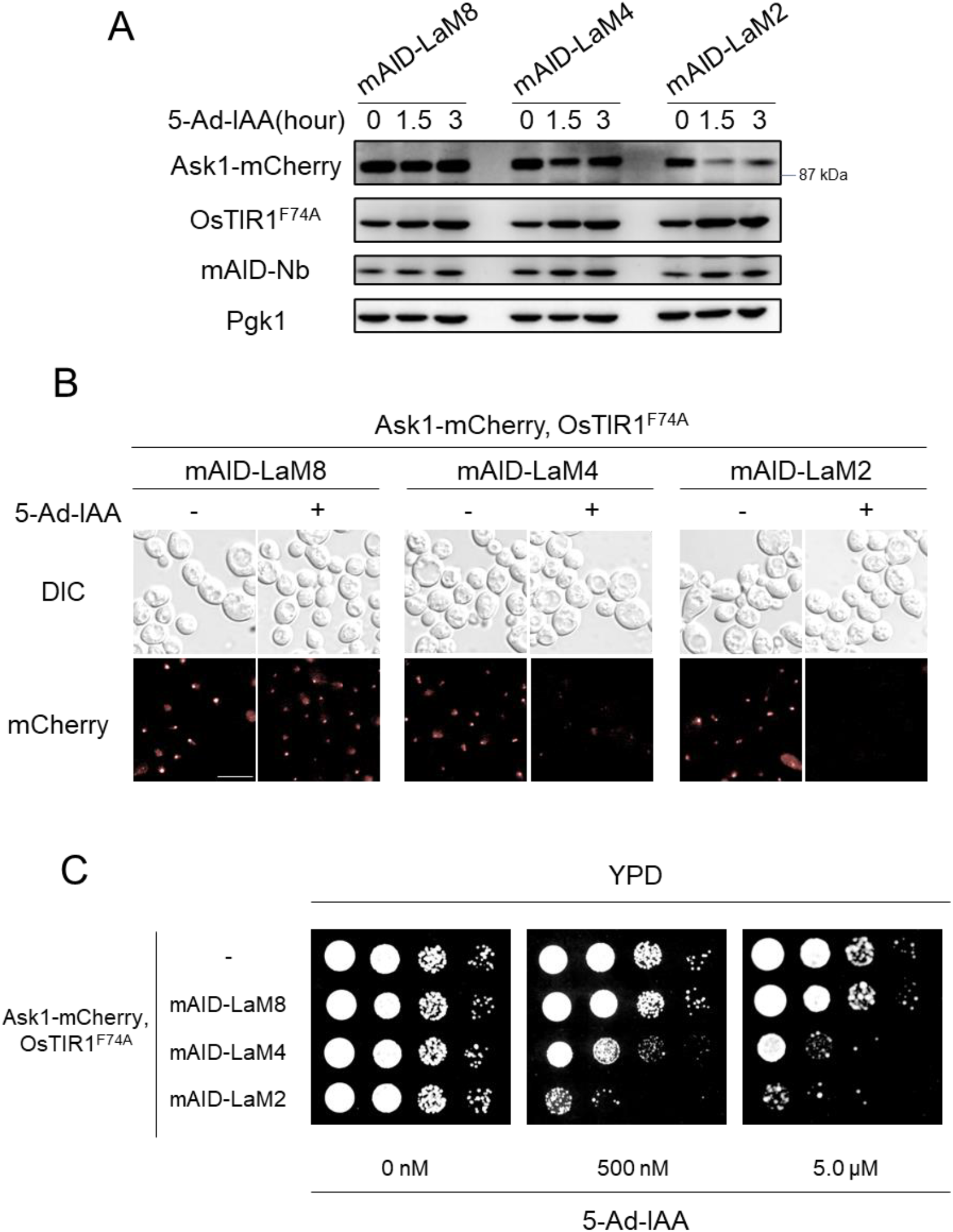
mCherry fusion protein is degraded effectively using LaM2 and LaM4. (A) Immunoblotting of endogenous tagged Ask1-mCherry. Cells expressing mAID-LaM8, mAID-LaM4, and mAID-LaM2 were established from the same cell line expressing OsTIR1^F74A^ and endogenous fusion Ask1-mCherry. (B) Fluorescence of Ask1-mCherry. Cells were treated with 5.0 μM 5-Ad-IAA for 3 h. Scale bar, 5 μm. (C) Serial dilution spotting on the YPD medium containing 5-Ad-IAA. Cells were grown for 24 h at 30 °C.

## DISCUSSION

Methods to induce target-protein degradation in cells often negatively affect the cells themselves. Inactivation of target proteins by temperature-sensitive mutants requires a temperature shift from 23–28 °C to 37 °C (35). Such severe temperature changes have significant effects on organisms. Furthermore, in conventional AID systems, target-protein degradation requires high concentrations of auxins, which negatively affects organisms. Our AlissAID system requires only 50 nM of 5-Ad-IAA for target-protein degradation, potentially preventing this problem.

In this study, we coincidentally found the levels of Neo1-mAID-GFP is reduced in the absence of 5-Ad-IAA in ssAID system. This protein reduction completely depends on the OsTIR1^F74A^ expression. Compared with OsTIR1^WT^, OsTIR1^F74A^ had a less sensitive to the natural auxin IAA but high concentration of IAA had an ability to induce the interaction OsTIR1^F74A^ and AID-tag at 30 °C in yeast two hybrid assay. Because this type of weak sensitivity to IAA was found in other mutants of OsTIR1 (OsTIR1^F74G^, OsTIR1^F74S^ or OsTIR1^F74C^) (11), TIR1 dependent basal degradation was found in some proteins with AID-tag in budding yeast. Interestingly, yeast two hybrid assay showed the sensitivity of OsTIR1^F74A^ to IAA was reduced at 37 °C, which might cause less basal degradation in mammalian cells. While basal degradation was not found in AlissAID system using VHHGFP4. Although 5-Ad-IAA directly induced interaction with OsTIR1^F74A^ and degradation target protein in ssAID system, 5-Ad-IAA induce the interaction with OsTIR1^F74A^ and degradation target protein indirectly in AlissAID system. This difference may produce mild basal degradation in AlissAID system in budding yeast. In conclusion, AlissAID system is useful to generate AID-based conditional knockdown cells with unstable protein in other AID systems.

Not all nanobodies (single-peptide antibody) work well in our AlissAID system. We tested three different anti-mCherry nanobodies (LaM2, LaM4, and LaM8). Among them, LaM2 induced target protein degradation most efficiently, whereas LaM4 weakly induced target degradation. In terms of mCherry-binding forces, LaM2 and LaM4 have KD values of 3.02 ± 1.88 nM and 22.50 ± 34.6 nM, respectively (36). Considering that the GFP-banding force of VHHGFP4 has a KD value of 1.40 nM (37), degradation efficiency appears to be associated with the affinity to the nanobody and target proteins. In terms of crystal structures, the mCherry LaM2 binding site is similar to the GFP binding site, which VHHGFP4 binds to (36,37). These binding sites may potentially be related to efficient ubiquitination and degradation. In conclusion, target-protein degradation efficiency is largely dependent on the type of nanobodies that are used in the AlissAID system.

In conclusion, this study demonstrates that GFP or mCherry fusion proteins can be degraded by the AlissAID system in budding yeast. In this study, we used nanobodies, which recognized GFP or mCherry proteins for degradation. Other binding domains, including monobodies (38) (39), DARPins (40) or binding peptides (41) that bind specific proteins can also be potentially used. The use of nanobodies, which recognize target protein directly in cells, would also enable degradation of untagged endogenous proteins using the AlissAID system. In budding yeast, there is a large clone collection of GFP-tagged strains. This clone collection enables the generation of various types of AlissAID strains systematically. A combination of the GFP clone collection and robotics would be useful for high-throughput screening and genome-wide analysis in budding yeast. Similar to the conventional AID system, we expect the AlissAID system to function in various eukaryotic organisms. This new method provides an ideal protein-knockdown system for a wide range of target proteins using various binding domains, including nanobodies.

## TABLE AND FIGURES LEGENDS

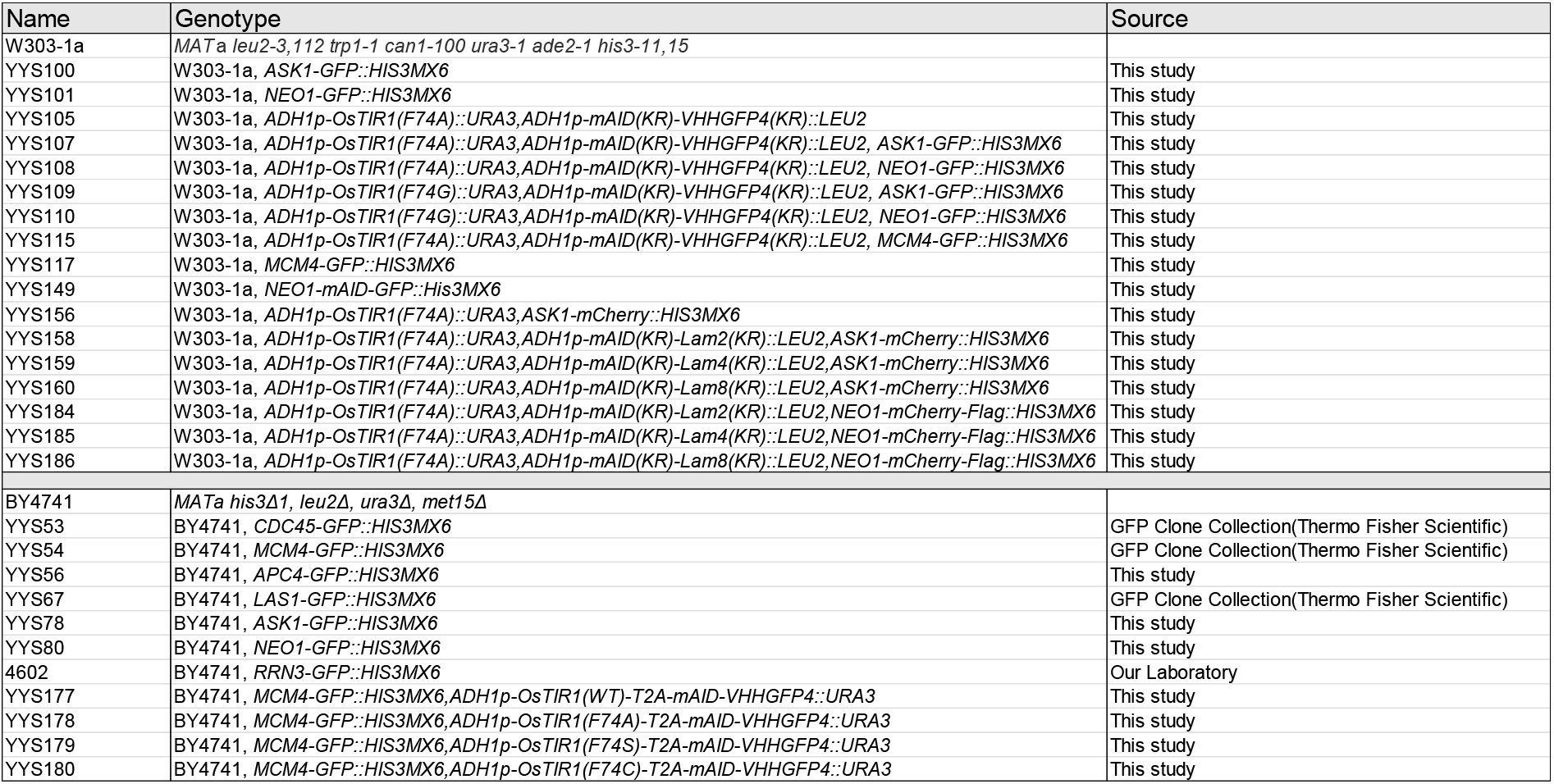

| Name | Genotype | Source |
| --- | --- | --- |
| W303-1a | *MATa leu2-3,112 trp1-1 can1-100 ura3-1 ade2-1 his3-11,15* |  |
| YYS100 | W303-1a, *ASK1-GFP::HIS3MX6* | This study |
| YYS101 | W303-1a, *NEO1-GFP::HIS3MX6* | This study |
| YYS105 | W303-1a, *ADH1p-OsTIR1(F74A)::URA3,ADH1p-mAID(KR)-VHHGFP4(KR)::LEU2* | This study |
| YYS107 | W303-1a, *ADH1p-OsTIR1(F74A)::URA3,ADH1p-mAID(KR)-VHHGFP4(KR)::LEU2, ASK1-GFP::HIS3MX6* | This study |
| YYS108 | W303-1a, *ADH1p-OsTIR1(F74A)::URA3,ADH1p-mAID(KR)-VHHGFP4(KR)::LEU2, NEO1-GFP::HIS3MX6* | This study |
| YYS109 | W303-1a, *ADH1p-OsTIR1(F74G)::URA3,ADH1p-mAID(KR)-VHHGFP4(KR)::LEU2, ASK1-GFP::HIS3MX6* | This study |
| YYS110 | W303-1a, *ADH1p-OsTIR1(F74G)::URA3,ADH1p-mAID(KR)-VHHGFP4(KR)::LEU2, NEO1-GFP::HIS3MX6* | This study |
| YYS115 | W303-1a, *ADH1p-OsTIR1(F74A)::URA3,ADH1p-mAID(KR)-VHHGFP4(KR)::LEU2, MCM4-GFP::HIS3MX6* | This study |
| YYS117 | W303-1a, *MCM4-GFP::HIS3MX6* | This study |
| YYS149 | W303-1a, *NEO1-mAID-GFP::His3MX6* | This study |
| YYS156 | W303-1a, *ADH1p-OsTIR1(F74A)::URA3,ASK1-mCherry::HIS3MX6* | This study |
| YYS158 | W303-1a, *ADH1p-OsTIR1(F74A)::URA3,ADH1p-mAID(KR)-Lam2(KR)::LEU2,ASK1-mCherry::HIS3MX6* | This study |
| YYS159 | W303-1a, *ADH1p-OsTIR1(F74A)::URA3,ADH1p-mAID(KR)-Lam4(KR)::LEU2,ASK1-mCherry::HIS3MX6* | This study |
| YYS160 | W303-1a, *ADH1p-OsTIR1(F74A)::URA3,ADH1p-mAID(KR)-Lam8(KR)::LEU2,ASK1-mCherry::HIS3MX6* | This study |
| YYS184 | W303-1a, *ADH1p-OsTIR1(F74A)::URA3,ADH1p-mAID(KR)-Lam2(KR)::LEU2,NEO1-mCherry-Flag::HIS3MX6* | This study |
| YYS185 | W303-1a, *ADH1p-OsTIR1(F74A)::URA3,ADH1p-mAID(KR)-Lam4(KR)::LEU2,NEO1-mCherry-Flag::HIS3MX6* | This study |
| YYS186 | W303-1a, *ADH1p-OsTIR1(F74A)::URA3,ADH1p-mAID(KR)-Lam8(KR)::LEU2,NEO1-mCherry-Flag::HIS3MX6* | This study |
| BY4741 | *MATa his3Δ1, leu2Δ, ura3Δ, met15Δ* |  |
| YYS53 | BY4741, *CDC45-GFP::HIS3MX6* | GFP Clone Collection(Thermo Fisher Scientific) |
| YYS54 | BY4741, *MCM4-GFP::HIS3MX6* | GFP Clone Collection(Thermo Fisher Scientific) |
| YYS56 | BY4741, *APC4-GFP::HIS3MX6* | This study |
| YYS67 | BY4741, *LAS1-GFP::HIS3MX6* | GFP Clone Collection(Thermo Fisher Scientific) |
| YYS78 | BY4741, *ASK1-GFP::HIS3MX6* | This study |
| YYS80 | BY4741, *NEO1-GFP::HIS3MX6* | This study |
| 4602 | BY4741, *RRN3-GFP::HIS3MX6* | Our Laboratory |
| YYS177 | BY4741, *MCM4-GFP::HIS3MX6,ADH1p-OsTIR1(WT)-T2A-mAID-VHHGFP4::URA3* | This study |
| YYS178 | BY4741, *MCM4-GFP::HIS3MX6,ADH1p-OsTIR1(F74A)-T2A-mAID-VHHGFP4::URA3* | This study |
| YYS179 | BY4741, *MCM4-GFP::HIS3MX6,ADH1p-OsTIR1(F74S)-T2A-mAID-VHHGFP4::URA3* | This study |
| YYS180 | BY4741, *MCM4-GFP::HIS3MX6,ADH1p-OsTIR1(F74C)-T2A-mAID-VHHGFP4::URA3* | This study |

