## Supplementary Figures for "Development of AlissAID system targeting GFP or mCherry fusion protein"

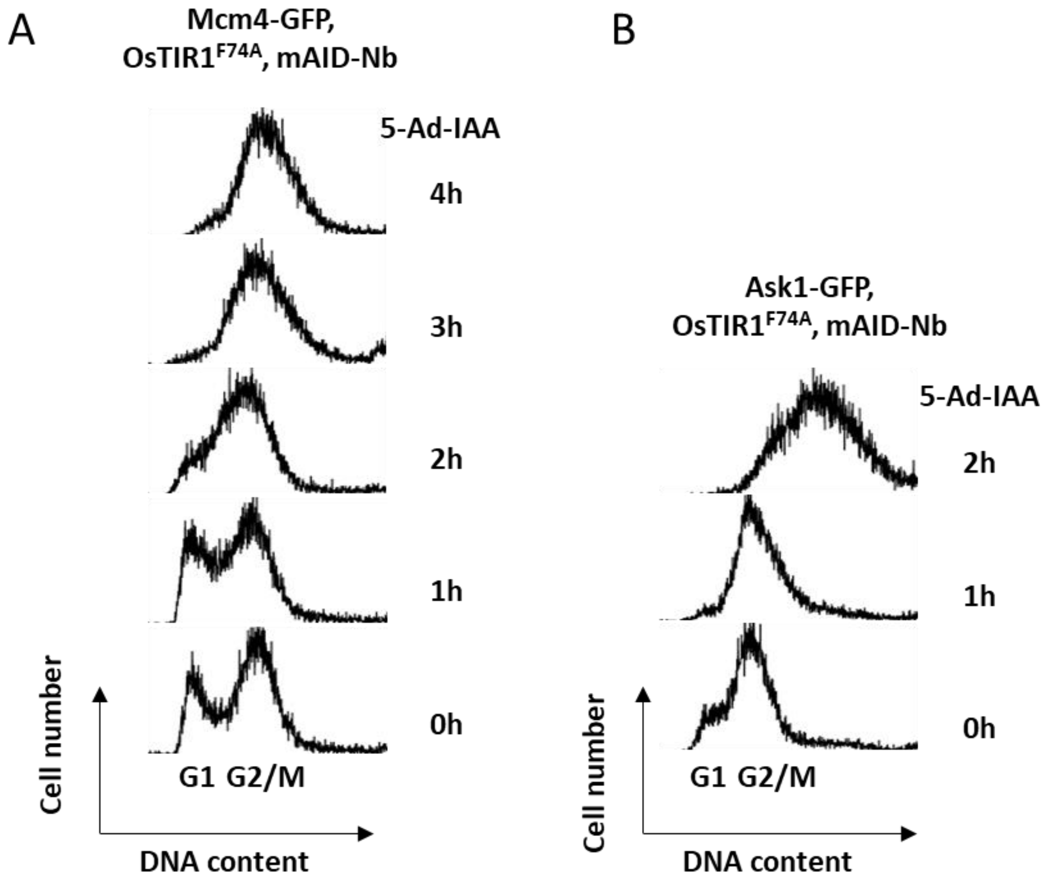


Supplementary Figure 1. (A) Cell cycle analysis of AlissAID strain targeting Mcm4-GFP. Cells were treated with 5 μM 5-Ad-IAA and sampled at the indicated time points. (B) Cell cycle analysis of AlissAID strain targeting Ask1-GFP. Cells were treated with 5 μM 5-Ad-IAA and sampled at the indicated time points.

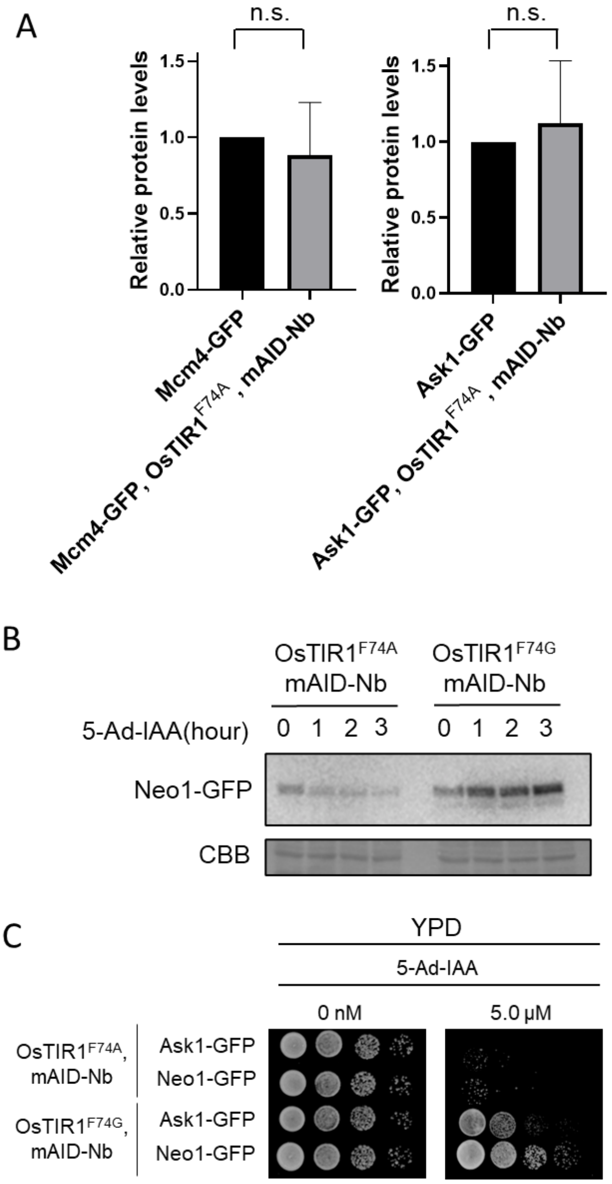


Supplementary Figure 2. (A) Immunoblotting of Mcm4-GFP and Ask1-GFP. Protein levels relative to OsTIR1$^{F74A}$ and mAID-Nb un-expression cell, normalized to those of Pgk1. (B) In OsTIR1$^{F74A}$ or OsTIR1$^{F74G}$ and mAID-Nb expression cells, Neo1-GFP degradation detected by Immunoblot analysis. Cells were treated with 5 uM of 5-Ad-IAA and samples are taken at indicated time points. CBB proteins are used as a loading control. (C) Serial dilution spotting on the YPD medium containing 5-Ad-IAA. Cells are grown for 24 h at 30 °C.

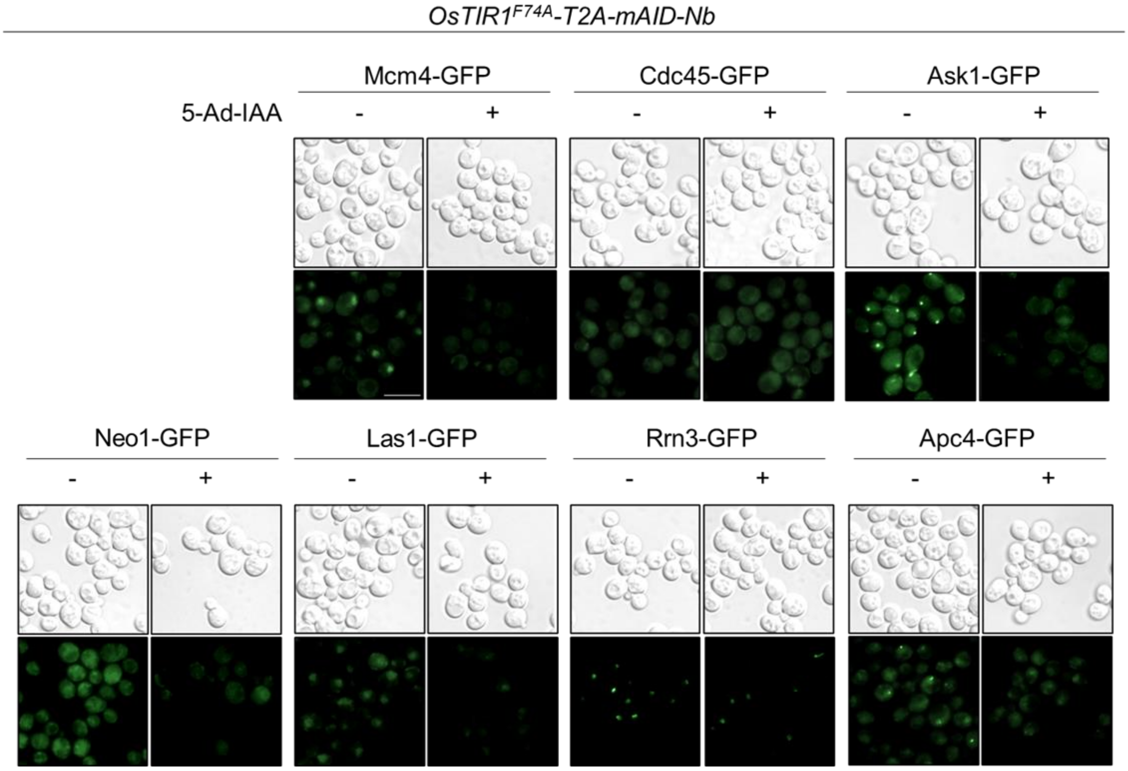


Supplementary Figure 3. Microscopic observation of various target proteins fused GFP. Cells were treated with or without 5 μM 5-Ad-IAA for 3 h. The white bar indicates 5 μm.

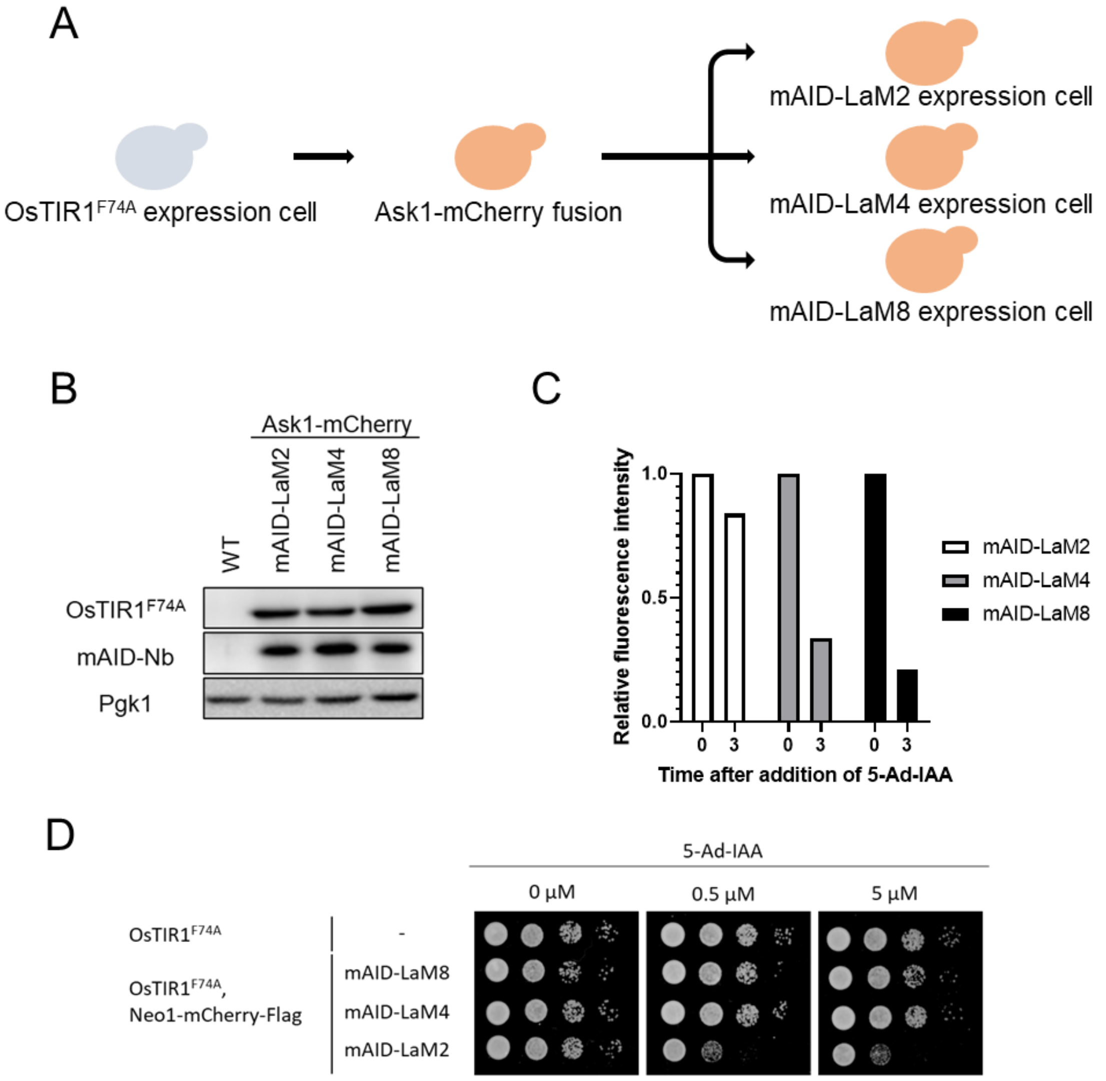


Supplementary Figure 4. (A) All LaM-expressing cell lines were established from mCherry tagged cell lines to the same OsTIR1$^{F74A}$-expressing line. (B) Immunoblotting of all LaM-expression cells absence of 5-Ad-IAA. (C) Fluorescence intensity of Ask1-mCherry in all LaM-expression cells. Cells were treated with or without 5 μM 5-Ad-IAA for 3 hours. Signals were measured using ImageJ and normalized with the number of cells in those visions. (D) Serial dilution spotting on the YPD medium containing 5-Ad-IAA. Cells are grown for 24 h at 30 °C.
